# Behavioral Interactions in Two Ant Species in The Southeast United States and Evidence for a Native Supercolony

**DOI:** 10.1101/2024.09.06.611643

**Authors:** Shreyas Kanwar, Renee Nowicki, Isabella R.E. Mavourneen, Joshua D. Gibson

## Abstract

Ants vary extensively in their colony structure, ranging from occupying single nests to tremendous supercolonies that occupy territories spanning large areas. Fewer than 1% of ants are known to produce supercolonies, yet they are disproportionately overrepresented in highly invasive ants. A broader understanding of supercolonial ants in their native range, therefore, may provide key insights into the factors that allow some ants to become invasive. Here, we show the results of behavioral assays of two native species of ants, *Dorymyrmex bureni* and *D. smithi* (Hymenoptera: Formicidae: Dolichoderinae) across geographically separated sites in southeastern Georgia, USA. We show that *D. smithi* has extremely low incidence of aggression in ants from sites up to 35 km apart, indicating that this may be a supercolonial species. In contrast, we show that *D. bureni* exhibits high levels of aggression between nest sites. *Dorymyrmex smithi* is also a temporary social parasite of *D. bureni* and these two species form mixed nests with no apparent interspecific aggression between workers, but we show that both species interact aggressively in assays within and between sites when the workers are derived from pure species nests. These findings represent an important addition to our knowledge of supercolonial species, and they also lay the groundwork for further studies of the parasitic relationship between these species.

## INTRODUCTION

Ants form colonies with a wide variety of social and nest structures, ranging from a single queen to many queens (monogynous or polygynous), and from residing in a single nest to spreading across many nests (monodomous or polydomous, Holldobler and Wilson 1977). In most ant species, regardless of the social and nest structure, the members of these colonies behave peacefully with one another and aggressively with members of other colonies from the same species (Bourke and Franks 1995). In polydomous colonies, this involves peaceful interactions between colony members from many spatially distributed nests while still maintaining aggressive interactions with ants from nests belonging to other colonies (Debout et al. 2007). Across the more than 150 polydomous ant species, there is tremendous variation in both the number of nests and the overall size of the colonies, ranging into up to thousands of nests and potentially forming supercolonies that can span up to several square kilometers in area (Debout et al. 2007, Helanterä 2022, Robinson 2014). Some invasive populations, for example Argentine ants (*Linepithema humile*), can form entire unicolonial populations in which all individuals interact peacefully regardless of the distance between their nests, including spanning continents (Helantera et al. 2009, van Wilgenburg et al. 2010). Given that supercoloniality is rare in ants in general but is prevalent in many invasive ants, a better understanding of supercolonial species that are currently confined to their native ranges is important to identifying the key factors that may lead to a species becoming invasive (Holldobler and Wilson 1977, Holway et al. 2002, McGlynn 2002). In studies of colony structure and identity, sensitive behavioral assays have been the key in identifying the limits of colony boundaries (Holldobler and Wilson 1990, Breed 2003). Therefore, we set out to use behavioral assays to better understand the colony structure of two understudied species of ants that are native to the Southeast United States, one of which produces large polydomous colonies.

*Dorymyrmex bureni* and *Dorymyrmex smithi*^*1*^ are both native ants in the Southeast United States, generally inhabiting the same disturbed sandy habitats, including sandhills, roadsides, and open fields (Nickerson 1976, Trager 1988, Whitcomb et al. 1972). These two ants are particularly interesting because they have strongly contrasting colony structures. *Dorymyrmex bureni* is a monodomous and monogynous species, whereas *D. smithi* is a polydomous and polygynous species (Nickerson et al. 1975b). The polydomy in *D. smithi* is rather extreme; Nickerson (1976) noted over 2,000 individual nests within a single field at Tall Timbers Research Station near Tallahassee Florida and regularly found several hundred nests clustered together. *Dorymyrmex smithi* has not been included in other descriptions of supercolonial ants, but this extensive polydomy is consistent with descriptions of supercolonies in other ants (Helanterä 2022). In addition to this stark contrast in colony structure, these ants also form a socially parasitic relationship in which *D. smithi* queens use *D. bureni* as a host colony during colony founding (Buren et al. 1975, Trager 1988). Mixed nests consisting of workers of both species have been observed in the field (though never directly intermixed with pure *D. smithi* nests), and when excavated they only contained a single *D. smithi* queen (Buren et al. 1975). These mixed nests were also identified in the field by Trager (1988) as well as in our own observations. Buren et al. (1975) found that these mixed nests did not persist for long periods of time and that they were eventually replaced by purely *D. smithi* nests, and the authors note that no incipient nests of purely *D. smithi* had been observed over several years of intense searching. Taken together, these findings strongly indicate that *D. smithi* queens use *D. bureni* as a host for new colony founding. Importantly, the researchers noted that workers of the two species from unmixed colonies reacted hostilely when they met in the field, with *D. smithi* workers typically holding their ground and flaring their mandibles and *D. bureni* workers quickly retreating. However, in the mixed colonies the two species’ workers seemed to be undertaking tasks equally and showed no hostility toward one another.

In this study, we examine the behavioral interactions of both *D. bureni* and *D. smithi* from three sites in southeastern Georgia that are located 35 km apart. We investigate aggressive behaviors in interactions between both species, as well as within nests and between sites for each species. In particular, we are interested in investigating whether *D. smithi* may form a large supercolony in this region. In addition, we investigate a specific mutual antennation behavior between *D. smithi* workers from within nests and between sites. These findings lay the groundwork for a better understanding of the social relationship between these two species and expand our knowledge of a potentially supercolonial ant species.

## MATERIALS AND METHODS

### Field collection

Worker ants of *D. bureni* and *D. smithi* were collected from May to June 2022 from three sites in southeastern Georgia: 1) Georgia Southern University’s Statesboro campus 2) the Statesboro S&S Greenway 3) George L. Smith State Park in Twin City. The two locations in Statesboro are ∼3 km apart, and these two are approximately 35 km from the state park location. Cheese puffs (CHEETOS^®^ Crunchy Cheese-Flavored Snacks, Frito-Lay, Plano TX) were used as bait to attract ants. Approximately 200 adult workers of each species were collected from each site using an aspirator and a 9-dram plastic vial (Bioquip, Rancho Dominguez, CA). No mixed nests were observed at these sites during worker collection, and all workers were collected from pure-species nests. *Dorymyrmex bureni* workers were collected from a single nest to ensure they were from the same monodomous colony and *D. smithi* were collected from the same small area, though due to their high nest density in their polydomous colony there were often multiple entrances within a few square meters (Nickerson 1976). Ants were transported back to the lab in plastic containers (Cut Comb Honey Container, 4 5/16″ x 4 5/16″ x 1 3/8″, 14 oz, Pioneer Plastics, Dixon, KY) with a lid with mesh vents and a coat of Insect-a-Slip (Bioquip, Rancho Dominguez, CA) on the walls to prevent ants from escaping.

### Laboratory conditions

Upon arriving in the lab, each box of ants was placed into its own 6-quart 13 5/8″ x 8 1/4″ x 4 7/8″ box (referred to as “nest box” henceforth), which also had a coat of Insect-a-Slip on the walls. The laboratory is maintained at approximately 25°C under a 12-12 light-dark cycle. Each nest box was provided with small nesting site consisting of a 100 x 15 mm petri dish that was painted black, had a layer of plaster in its base that was sprayed with water prior to adding it to the nest box, and with a small entrance hole drilled into the side. Water and a 20% sugar/water solution were provided in cotton-plugged test tubes, which were replaced as needed. During their time in the lab, ants were provided with <5 g of scrambled egg 2-3 times per week to allow *ad libitum* feeding.

### Behavior assays

We performed five types of comparisons in our behavioral assays and each type included all possible pairwise comparisons of sites: between species (three sites per species yielded nine pairwise comparisons), within nest boxes for each species (three pairwise comparisons per species), and between sites for each species (three pairwise comparisons per species). All pairwise comparisons were replicated 12 times, yielding 108 between-species assays and 36 assays for all other comparison. Ants for behavioral assays were placed into an Insect-a-Slip lined 100 x 15 mm petri dish along with a small damp cotton ball and were allowed to acclimate in this dish for 24 hours prior to the assays. Separate petri dishes were used for each nest box, and each contained 25 ants. If more than 20 ants were needed for the behavioral assays, an additional petri dish containing 25 ants was set up to ensure all ants experienced the same conditions prior to behavioral assays. All behavioral assays were conducted one-on-one in an Insect-a-Slip lined well of a 12-well cell culture plate (Greiner Bio One, Monroe NC), and up to six assays were run simultaneously in the same plate. Individual ants were only used for a single assay and were not placed back in their nest box. All assays were conducted for 10 minutes and were recorded using a Canon EOS 6D Mark 2 dSLR camera with a Canon 100 mm f/2.8L Macro lens (resolution: 1080p). For each behavioral assay, one ant from each pair were placed in a well of one of two culture plates using an aspirator. These plates were attached with tape on their long edge so the “top” plate could be quickly tipped over, depositing each ant into the well of the “bottom” plate containing its corresponding assay partner. Prior to tipping the ants over, they were allowed to recover from aspiration for approximately 5 minutes. The behavioral assay and video recording was initiated immediately after tipping the top plate over to ensure that all behaviors were recorded. A spreadsheet was used to randomly assign each assay to a plate and well, and ants were split evenly between the top and bottom plates for comparisons across sites/species to account for any behavioral impact of being tipped into another well.

### Aggression scoring

Behavioral assays were scored blindly using the video footage and each assay was scored separately by two reviewers; while between-species assays were apparent due to the ant’s coloration, the scorer did not know which sites the ants were from, nor did they know if they were viewing within-nest or between-site assays for each species. A four-point scale of aggression was used (modified from Suarez et al. 1999, Suarez et al. 2002): 0 = no contact; 1 = antennation; 2 = non-contact aggression, mandible flaring, gaster curling; 3 = biting, nipping; 4 = grappling, tumbling, prolonged fighting and/or killing. Each 10-minute assay was assigned the score of the highest level of aggression observed, though no scores were assigned in the first 10 seconds after tipping the top plate over. Assays were binned into non-aggressive (score of 1) or aggressive (score of 2, 3 or 4), but were also placed into bins based on whether the ants physically touched one another aggressively (score of 3 or 4) or not (score of 1 or 2). We refer to these two types of aggression as non-contact aggression (score of 2) or contact aggression (score of 3 or 4). This latter strategy was employed because contact aggression such as physical biting and grappling are less ambiguous than mandible flaring and gaster curling.

### Antennation scoring

In the within-nest and between-site comparisons of *D. smithi*, we also separately scored a behavior that we term “mutual antennation.” This behavior involves two ants facing one another and quickly and intensely antennating each other. When this antennation was observed, we scored the assays based on their duration; short (2-5 seconds, never longer, in at least two bouts), intermediate (>5 seconds in at least one bout, for a total time of between 10 and 30 seconds during the assay), and extended (>5 seconds in at least one bout, total time of >30 seconds during the assay). Two of the within-nest assays could not be scored for this behavior because the ants were on the wall of the well and we couldn’t observe their antennae.

### Statistical Analysis

Pairwise comparisons of all proportions were tested in 2×2 contingency tables using Fisher’s Exact Test (JMP®, Version 17.0.0 SAS Institute Inc., Cary, NC) and the Holm-Bonferroni method was applied to correct for multiple testing (Holm 1979). To compare aggression frequency, we tested all pairwise combinations of our five assay types except for the two within-nest comparisons (due to both having zero aggressive interactions), for a total of nine pairwise tests (corrected *p*-value of 0.005). Pairwise comparisons of within-nest and between-site frequency of antennation were tested for each of the three duration categories (corrected *p*-value of 0.017).

## RESULTS

### Field observations

Each of the three sites was consistent with habitat described in previous literature, they generally have sparse vegetation and are near disturbed areas (Nickerson 1976, Trager 1988, Whitcomb et al. 1972). The Georgia Southern

University campus site is on an open sandhill bordering a paved road, the S&S Greenway site is in a disturbed area with little vegetation next to a paved path and roadway, and the George L. Smith site is on a sandhill next to a dirt trail. We did not explicitly measure nest distribution, but our observations match those of previous descriptions (Buren et al. 1975, Nickerson et al. 1975b). The *D. smithi* colonies were extensive, spanning tens of meters at each site, though we did not measure the furthest nest sites or their area to determine maximum colony size. Nests of *D. bureni* and *D. smithi* were not intermixed, but *D. bureni* nests were found on the periphery of the *D. smithi* colony. We did not observe any mixed nests at these sites during the collection period.

### Aggression assays

In all assays of aggression, non-contact aggression without escalating to physical contact-aggression was rare and did not significantly alter the proportion of aggression for any assay nor did it change any significance of comparisons between assay types (Fig. 1). Therefore, we used the proportion of physical aggression for subsequent comparisons. Interspecific aggression assays from all nine pairwise comparisons indicated high frequencies of contact aggression (Fig. 1). The frequency of aggression between sites of *D. bureni* was similarly high (Fisher’s Exact Test, FET, *p* = 0.4044) and was significantly higher than the frequency of aggression observed in within-nest assays (FET, *p* < 0.0001). However, the frequency of aggression between sites of *D. smithi* was significantly lower than the frequency between species and the frequency between *D. bureni* nests (FET, *p* < 0.0001 for each), and it was not significantly higher than the complete lack of aggression observed in within-nest assays of *D. smithi* (FET, *p* = 0.2394).

**Fig 1.**
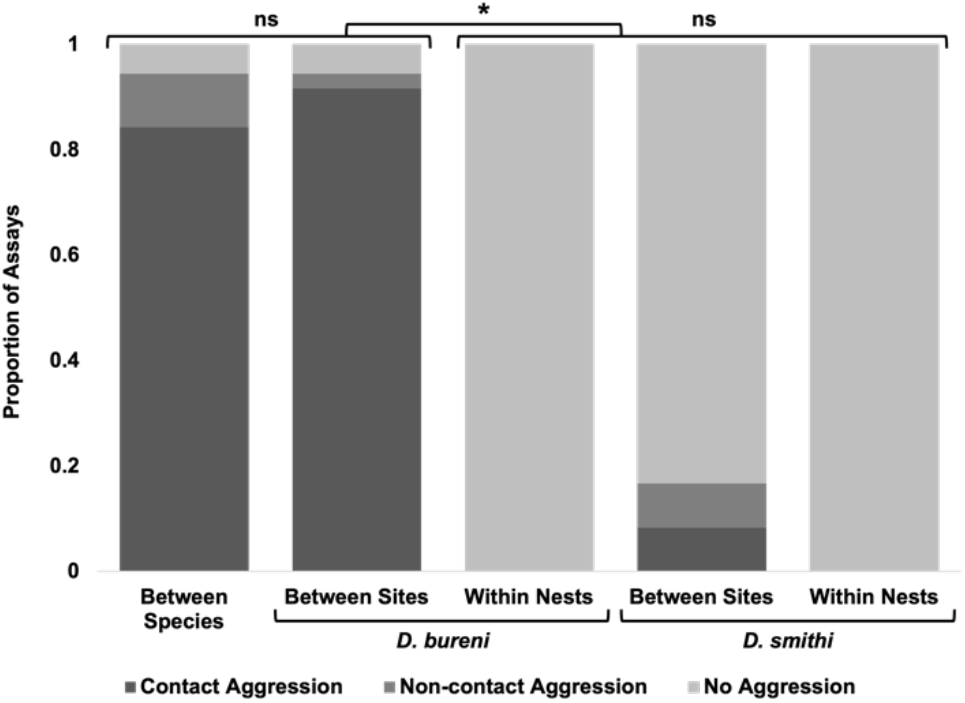
Proportion of behavioral assays in each aggression category across comparison types. Top bars indicate significance for all pairwise comparisons except between the two within-nest categories, which were not tested due to zero observations of aggression. Significance was corrected using the Holm-Bonferroni method. Significant differences are indicated with * and non-significant differences are indicated with ns

### Antennation

The frequency of *D. smithi* assays with short or intermediate duration “mutual antennation” did not differ when between-site and within-nest assays were compared (Fig. 2; short, FET, *p* = 0.2999; intermediate, FET, *p* = 0.5743). However, extended duration antennation occurred over six times more frequently in between-site assays than within-nest assays (FET, *p* < 0.0001). A table summarizing both the aggression scores and the antennation observations for each category of assay is provided in Online Resource 1.

**Fig. 2.**
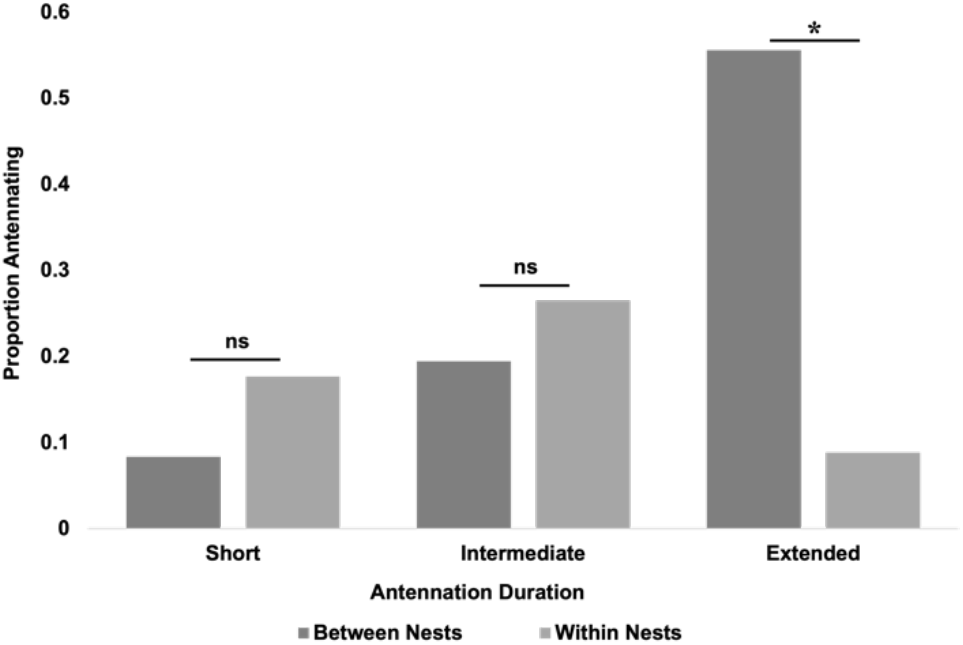
Proportion of assays in each antennation duration category in between-site and within-nest assays of *Dorymyrmex smithi*. Top bars represent significance of pairwise comparisons. Significance was corrected using the Holm-Bonferroni method. Significant differences are indicated with * and non-significant differences are indicated with ns

## DISCUSSION

Our results are consistent with previous studies indicating that *D. bureni* is a monodomous species and that *D. smithi* is a polydomous species, while also indicating that *D. smithi* has the characteristics of a supercolonial species like many invasive ants. Our results also support the anecdotal evidence that, while workers of these two species cooperate when in mixed parasitic nests, they exhibit high levels of aggression with one another when they come from nests composed solely of their own species.

Our observations in the field support previous descriptions of these ants’ nesting strategies (Buren et al. 1975, Nickerson et al. 1975b). *Dorymyrmex bureni* workers were only observed entering and exiting individual nests while *D. smithi* workers were generally dispersed across the surface of the entire site and there was no indication of nest fidelity. We did not explicitly map out the locations of nests, but the *D. bureni* nests were typically located outside of the “territory” of the *D. smithi* colony (i.e. they were not interspersed with the *D. smithi* nests). Each site had many *D. smithi* nests that were located across a broad area. There was no indication of any aggression between *D. smithi* at any location. We did not observe interactions between *D. bureni* from different nests nor between the two species in the field.

In our laboratory assays, we observed a very high frequency of aggression in all between-site assays of *D. bureni* while never observing aggressive interactions when comparisons were made with nestmates (Fig. 1). Given that these ants were living peacefully in the same nest box prior to the assays, this latter finding indicates that our assay conditions don’t induce aggressive interactions in what would otherwise be peaceful interactions. Similarly, we observed a very high frequency of aggression in all assays between the two species. Ants often display aggressive behavior to different species, though this depends on the species being compared and can also be context dependent (Chong and Lee 2010, Human and Gordon 1996, Lemoli et al. 1984, Neumann and Pinter-Wollman 2022, Tanner and Adler 2009). We observed this high frequency of interspecific aggression regardless of whether each species came from the same site or different sites, indicating minimally that locality does not appear to influence interspecific aggression in these species. This is important because previous observations, as well as our own in the field (Nowicki, personal observation), show that workers within mixed colonies of these two species do not undergo this kind of aggressive interspecific interaction (Buren et al. 1975). This indicates that there is a clear behavioral shift when these ants are living in mixed, parasitic nests, though the mechanism underlying this shift is unknown.

We never observed aggressive interactions in any within-nest interactions of *D. smithi* (Fig. 1). As with the identical result in *D. bureni*, this is expected because these ants were collected from the same nests and maintained together in the same nest boxes in the lab, but it further supports that our assays do not induce aggressive interactions. Interestingly, aggressive interactions in between-site comparisons of *D. smithi* were very rare, including between sites that are approximately 35 km apart (Fig. 1). Moreover, in the few instances of aggression, the intensity of the aggression was relatively low. In the three instances of physical aggression (of 36 assays), only one involved prolonged fighting designated as a four on our aggression scale (see Online Resource 1 for a summary of assay scores). This contrasts with the high proportion of assays that involved prolonged fighting in both the interspecific assays (57 of 108 assays) and the between-site assays of *D. bureni* (30 of 36 assays). This lack of aggression across a broad geographic range indicates that *D. smithi*, in addition to being polydomous at each site, may be a rare example of a native supercolonial ant (Helanterä 2022). Interestingly, while the between-site assays lacked aggression, they did exhibit extended bouts of antennation over six times more frequently than the within-site assays (Fig. 2). A study of another native supercolonial species, *Formica paralugubris* in Switzerland, similarly found increased levels of antennation in nonnestmates in populations showing otherwise low levels of aggression (Holzer et al. 2006). These findings indicate that *D. smithi* recognizes nonnestmates, though it is not clear if this longer antennation indicates differences in colony membership between sites or if it simply indicates that these ants spend additional time assessing one another when they are unfamiliar. We did not run assays with *D. smithi* collected from different nests at the same site or that were kept in different nestboxes in the lab, so we cannot assess how these ants interact with ants from different nests that we know belong to the same colony. The antennation behavior we noted was always a close head-to-head process and did not include bodily antennation. In many instances it appeared that the ants may be engaging in, or attempting to engage in, the exchange of food via trophallaxis due to one or both slowly opening their mandibles while antennating and at times appearing to touch mouthparts. However, we were unable to consistently identify whether this was trophallaxis based on the quality of the video recordings. Ants have been shown to use trophallaxis for a wide range of purposes, including for both colony social maintenance via homogenization of colony odor, and as a form of appeasement when aggression is present (Meurville and LeBoeuf 2021). If this behavior in *D. smithi* does involve trophallaxis, the lack of any other aggression may indicate that this is a process of attempting to unify the colony odor across the supercolony, though further study is needed to disentangle the many possibilities.

There is no indication that *D. smithi* has been introduced or is invasive anywhere, but in addition to its apparent supercolonial colony structure it shares other characteristics with highly invasive ants. It is found in naturally disturbed areas, similar to many invasive ant species (Holway et al. 2002, Trager 1988) It is aggressive and ecologically dominant over other ants when found together, predating 97% of founding queens of the highly invasive red imported fire ant, *Solenopsis invicta* (Nickerson et al. 1975a). It has also been shown to tend to and protect at least 27 species of plant-feeding Homopteran insects (Lach 2003, Styrsky and Eubanks 2007, Nickerson and Whitcomb 1988). Another recently described native supercolonial species in Ethiopia, *Lepisiota canescens*, appears to have invasive characteristics as well but also has not been found outside its native range (Sorger et al. 2017). In both cases, this supercolonial structure and their presence in disturbed habitats may indicate a potential to become invasive if introduced outside their native range.

We didn’t specifically investigate the parasitic relationship of these two species, it is an important factor to consider when assessing whether this supercolonial species could become invasive. The parasitic lifestyle of *D. smithi* would seem to be a major factor limiting its ability to be introduced and become established, as it could require that its host species, *D. bureni*, be present in the introduced range as well. This parasitic relationship hasn’t been extensively studied, but our current understanding points to one of obligate temporary social parasitism during new colony founding by *D. smithi*. The strongest evidence for this being obligate parasitism is that of Buren et al. (1975), who were unable to identify any newly formed incipient nests *D. smithi* in Florida despite several years of intensive surveying, while they identified several incipient nests of *D. bureni*. Currently, the ontogeny of a newly established *D. smithi* nest expanding into a colony containing hundreds of nests is unknown, but it appears that existing *D. smithi* nests can establish additional nests either with or without parasitism. Trager (1988) notes that on two occasions, known *D. bureni* nests on the periphery of *D. smithi* colonies became mixed nests over the course of several days, indicating that *D. smithi* workers can invade neighboring *D. bureni* nests. However, the density of nests within a *D. smithi* colony are very high (2-4 nests per square meter) and these densities are not observed for the monodomous *D. bureni* (Nickerson 1976), indicating that *D. smithi* workers are very likely able to build new nests while expanding from an existing nest without the need for *D. bureni* as a host. Taken together, these findings indicate that *D. smithi* queens are unable to establish a new colony without parasitism, but existing nests can spread through a budding process without the need for the host species. Despite obligate temporary social parasitism, this still may indicate that *D. smithi* has the potential to become established as an expanding supercolony. This is very similar to the highly invasive Argentine ant, *Linepithema humile*, despite them not being a social parasite. Argentine ant queens do not perform mating flights and are unable to found colonies without existing workers (Hee et al. 2000, Passera and Keller 1990). Instead, colonies naturally spread by budding, and propagules of workers, brood, and queens that are transplanted by humans result in these ants spreading throughout the invasive range (Suarez et al. 2001). If the other aspects of *D. smithi* natural history are conducive to thriving in an introduced area, then their need for a host species to establish new colonies may not be a significant barrier to becoming invasive.

We have shown that *D. smithi* appears to form supercolonies with virtually no aggression across a broad geographic span while it maintains high levels of aggression with *D. bureni* when derived from a non-mixed nest environment. We have also shown that *D. smithi* shows recognition of nonnestmates through increased mutual antennation, though it is unclear if this behavior may indicate very subtle aggression or is a mechanism of maintaining colony unity. Further work is needed to assess additional sites at increased distances to determine if there may be multiple supercolonies of *D. smithi* or if they may represent a unicolonial population. Finally, this study sets the groundwork for future studies to investigate the interactions of these species in their parasitic relationship.

## Supporting information

Online Resource 1 (table)

## ACKNOWLEDGMENTS

The authors would like to thank Lisa D. Brown for feedback on the manuscript. This project received funding from the National Science Foundation Research Experiences for Undergraduates awards (DBI 1757536 and 2244232) made to Georgia Southern University to support SK and IM, and a Summer Graduate Student Research Assistantship from the James H. Oliver, Jr. Institute for Coastal Plain Science at Georgia Southern University to partially support RN.

## Footnotes

1 Ants referred to as *D. smithi* currently range across the southern US, from Florida to Utah, however there is some dispute over whether this is a single species (see Trager 1988, *Conomyrma medeis*; Snelling 1995 *D. smithi*). Here we use the most current name (Snelling 1995) and note that all citations for biological data are derived from the same southeastern populations/species that we have studied.

