## Supplementary material for "Behavioral Interactions in Two Ant Species in The Southeast United States and Evidence for a Native Supercolony": Online Resource 1 (table)

| | Number of assays in each category of aggression | | | | | | | Number of assays in each category of annentation duration | | | | | |
|---|---|---|---|---|---|---|---|---|---|---|---|---|---|
| **Comparison type** | **1** | **2** | **3** | **4** | **Total assays** | **Total all aggression (scores 2-4)** | **Total contact aggression (scores 3-4)** | **No antennation** | **Short** | **Intermediate** | **Extended** | **Total antennating** | **Total assays** |
| Between species | 6 | 11 | 34 | 57 | 108 | 102 | 91 | | | | | | |
| Within bureni nests | 36 | 0 | 0 | 0 | 36 | 0 | 0 | | | | | | |
| Within smithi nests | 36 | 0 | 0 | 0 | 36 | 0 | 0 | 16 | 6 | 9 | 3 | 18 | 34 |
| Between bureni nests | 2 | 1 | 3 | 30 | 36 | 34 | 33 | | | | | | |
| Between smithi nests | 30 | 3 | 2 | 1 | 36 | 6 | 3 | 6 | 3 | 7 | 20 | 30 | 36 |
